# Finding signatures of low-dimensional geometric landscapes in high-dimensional cell fate transitions

**DOI:** 10.1101/2025.05.27.656406

**Authors:** Maria Yampolskaya, Laertis Ikonomou, Pankaj Mehta

## Abstract

Multicellular organisms develop a wide variety of highly-specialized cell types. The consistency and robustness of developmental cell fate trajectories suggests that complex gene regulatory networks effectively act as low-dimensional cell fate landscapes. A complementary set of works draws on the theory of dynamical systems to argue that cell fate transitions can be categorized into universal decision-making classes. However, the theory connecting geometric landscapes and decision-making classes to high-dimensional gene expression space is still in its infancy. Here, we introduce a phenomenological model that allows us to identify gene expression signatures of decision-making classes from single-cell RNA-sequencing time-series data. Our model combines low-dimensional gradient-like dynamical systems and high-dimensional Hopfield networks to capture the interplay between cell fate, gene expression, and signaling pathways. We apply our model to the maturation of alveolar cells in mouse lungs to show that the transient appearance of a mixed alveolar type 1/type 2 state suggests the triple cusp decision-making class. We also analyze lineage-tracing data on hematopoetic differentiation and show that bipotent neutrophil-monocyte progenitors likely undergo a heteroclinic flip bifurcation. Our results suggest it is possible to identify universal decision-making classes for cell fate transitions directly from data.

## I. INTRODUCTION

Many animals have complex organs with specialized cells that coordinate to maintain homeostasis. Cells differentiate into these cell types over the course of embryonic development. In each cell, thousands of genes work together and receive information from chemical signals and mechanical cues to determine the cell’s lineage and function [1]. Although gene regulation is complex and high-dimensional, cells follow remarkably consistent developmental programs [2, 3]. The ways cells transition between types is robust: even when perturbed, they tend towards these development trajectories. Differentiated cells are stable in their cell fate unless exposed to particular signals. In injury and in experimental protocols specially designed to induce transdifferentiation, even mature adult cells can change their type [4–7]. An open problem in cell biology is understanding how cells retain and transition between cell fates in development, injury, and directed differentiation.

The experimental approach to understanding cell fate transitions often involves probing a cell’s gene expression profile [8]. Gene expression is a major factor in determining which proteins are created, so observing gene expression gives information about a cell’s function and therefore its type. Single-cell RNA-sequencing (scRNA-seq) is a widely-used technique for measuring gene expression profiles by capturing RNA from individual cells [9, 10]. There is an abundance of scRNA-seq data describing cell type, often in the form of scRNA-seq atlases that span entire organisms [11–13]. Despite the quantity of data, the theory of cell fate transitions is still in its infancy [14].

One of the earliest theoretical efforts to understand these transitions is the Waddington landscape, a metaphor for conceptualizing differentiation [15]. Because cell types are robust and discrete, Waddington proposed that cell fates act like attracting basins in a landscape. The developing cell is represented by a ball rolling down valleys to end up in one such basin. This metaphor suggests that the complex process of differentiation is effectively a low-dimensional landscape. The formalization of the Waddington landscape has been a long-standing problem, often approached from the perspective of dynamical systems [16–22]. In the language of dynamical systems, the Waddington landscape says that cell types act like attractor states: stable steady states which draw in nearby points.

Building on this work, Rand et al. [23] argue that we can categorize cell fate decisions and their landscapes according to general qualitative classes defined by bifurcations. These classes are characterized by the numbers of attractors, saddle points (which, in three dimensions, can be thought of as hills between attractor basins) and the paths connecting these objects. These classes are generic: any dynamical system with an associated landscape and attractor states can be described at least locally by these decision-making classes. Identifying classes of transitions is powerful because it opens the door to finding common, universal features across transitions, a deeper understanding of how cell fates relate to one another, and the ability to predict transitions based on general principles.

Although the landscape framework has been applied to a variety of biological contexts [21, 23– 25], the connection between cell fate landscapes and gene expression is not well-defined. Since gene expression is the primary way to experimentally probe cell fate, it’s important to bridge this gap. To unify the spaces of cell fate and gene expression, previous work has modeled gene regulatory networks as Hopfield networks: high-dimensional networks where particular stored patterns act as attractor states [26–36]. In this context, the stored patterns correspond to the gene expression profiles of cell fates. This description allows one to define cell fate coordinates calculated from gene expression data. These Hopfield-inspired coordinates have been applied to both bulk and single-cell RNA-seq data.

However, Hopfield models typically have static attractors whereas actual cell fate transitions involve signals which destabilize attractors, causing bifurcations which can be described with land-scapes and decision-making classes. In this paper, we introduce a model that combines the pattern retrieval properties of modern Hopfield networks with decision-making classes. This model captures the essential spaces of cell fate decisions: the dynamics occur in gene expression space while traversing a cell fate landscape that is controlled by received signals. The model predicts the cell fate trajectories corresponding to different landscapes and classes of decision-making. These cell fate trajectories are drawn on cell fate coordinates that can be calculated directly from scRNA-seq data, making it possible to directly compare candidate landscapes to experimental data. This allows us to identify signatures of generic decision-making classes and search for these signatures in experimental time-series data.

To illustrate the interpretative power of our model, we predict the dynamics of differentiation corresponding to three classes of decision-making and identify signatures of each one. To search for these signatures in experiment, we calculate cell fate coordinates from time-series scRNA-seq data. In a dataset describing the maturation of mouse lung epithelial cells, we find evidence of a triple cusp class. In an analysis of mouse hematopoietic lineage tracing data, we see trajectories suggestive of double cusp and heteroclinic flip classes. These studies show that our model, combined with time series scRNA-seq data, can directly compare general decision-making classes to real transitions, revealing universal properties and relationships between cell fates.

## II. RESULTS

### A. A model of differentiation unifying the spaces of gene expression, cell fate, and signaling

There are three mathematical spaces relevant to cell fate transitions: gene expression, cell fate, and signaling. We introduce a model which captures dynamics across the three relevant spaces. Using gene expression, we define cell fate coordinates to make the abstract cell fate landscape concrete and quantifiable. Our model describes cell fate dynamics by defining a landscape on these coordinates, with attractor basins determined by signals. Figure 1 illustrates the structure of the model. It is agnostic to the particular biological context. This is a universal framework that can be applied to cell fate transitions in development, injury, and directed differentiation. The basic assumption is that a cell type is a measure of cell identity that is stable under certain environmental and signaling conditions. Even transient attractors, like temporary progenitor cell types in development that eventually lead to mature types, can be captured by this model given that they are stable under some conditions.

**FIG. 1:**
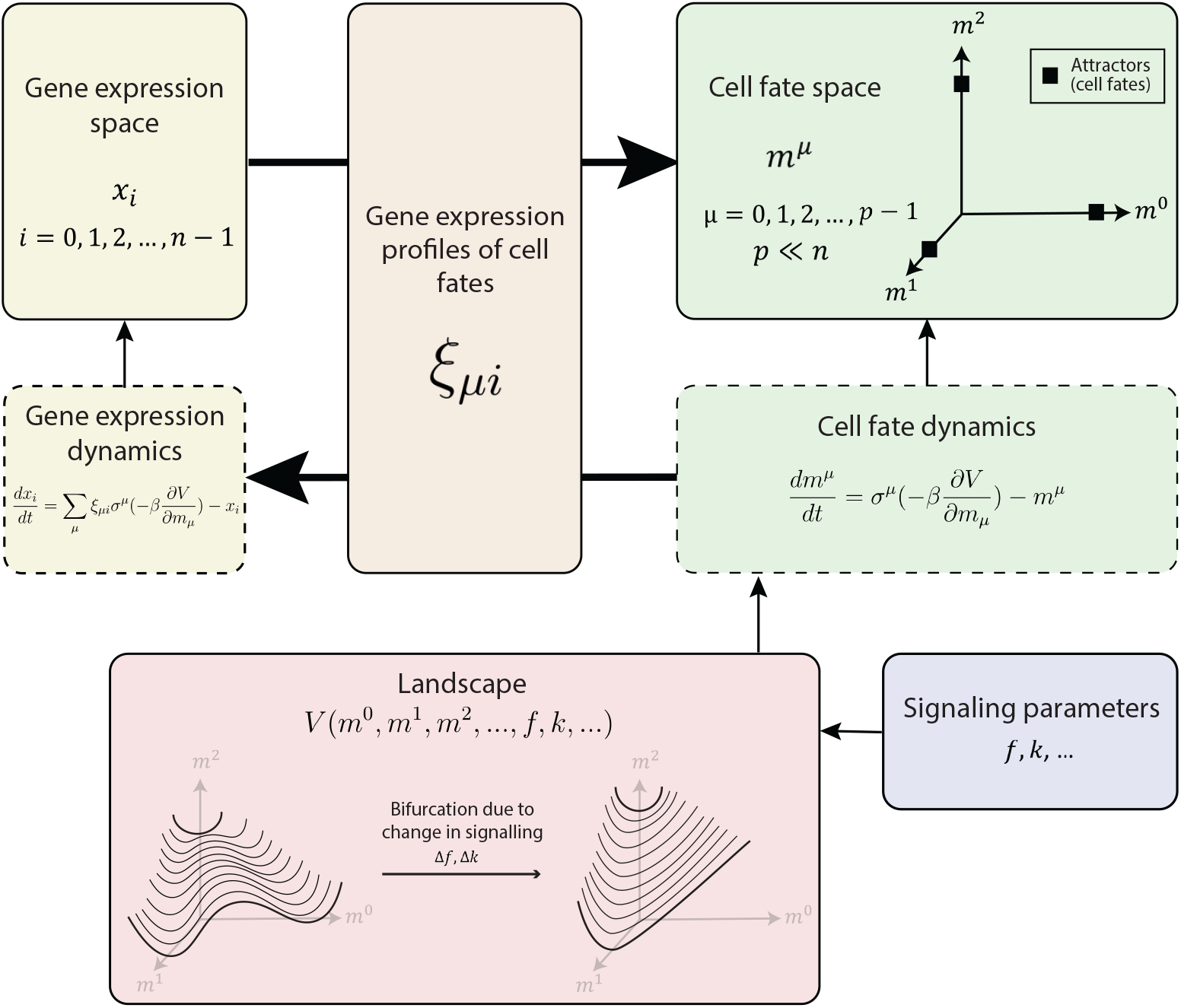
A model of cell fate transitions translates between high-dimensional gene expression space and a low-dimensional landscape. The model represents the gene activity and cell fate of one cell, with the gene expression of the *i*th gene in the cell indicated by *x*_*i*_. The Hopfield-inspired order parameter *m*^*µ*^ measures the alignment of the system with cell fate *µ* using the gene expression profiles of cell fates (*ξ*_*µi*_, which is the gene expression of gene *i* in cell fate *µ*). The order parameters *m*^*µ*^ form a p-dimensional cell fate space, where *p* is the number of cell fates. The *µ*th cell fate corresponds to the point *m*^*µ*^ = 1, *m*_≠*µ*_ = 0. The dynamics of *x*_*i*_ and *m*^*µ*^ have the pattern retrieval behavior of the Hopfield model, where cell fates act as attractors, while also traversing an input cell fate landscape *V*. *V* depends on signalling parameters *f, k*, … such that changes in signals received by the cell are represented by changes in these parameters, and these changes may cause a bifurcation in the landscape.

Our mathematical construction is based on a generalization of the Modern Hopfield network, a dynamical system for associative memory [26, 27, 37]. Building on previous work, we exploit the close relationship between recurrent neural networks for associative memory and Waddington’s landscape [29, 31, 36]. In both of these systems, the dynamics are defined in terms of an interacting network whose attractors we wish to specify. In the Hopfield model, the nodes of the network correspond to the firing rate of neurons and the attractors to stored memories. In the context of development, the nodes of our network correspond to the expression of different genes and the attractors to developmentally stable cell fates.

Our model builds upon Modern Hopfield networks by implementing two important changes that allow us to model cell fate transitions: (1) the use of generalized order parameters and (2) the inclusion of signal-dependent forces that cause transitions between attractors [38]. With these changes, the attractors (or memories) in our network change stability in response to external cues via signal-induced bifurcations. Additionally, the dynamics of the modified model do *not* have a Lyapunov function and *cannot* be described purely in terms of gradients of a potential function.

Nonetheless, many of the qualitative dynamical properties can still be understood in terms of landscapes. In the main text, we focus on giving an overview of our model. A detailed discussion of our construction can be found in the Methods and Supplemental Information, including a discussion of the relationship between this model and pure gradient systems.

In our model, the state of a cell is specified by its gene expression profile. At any time *t*, the state of the system is described by a *N*-dimensional real-valued vector *x*_*i*_(*t*) that encodes the expression of gene *i* at time *t*. We focus on the differentiation dynamics of a system with *p* possible cell fates (*µ* = 0, …, *p −* 1). These cell fates are defined by a list of *p* vectors in gene expression space, {*ξ*_*µi*_}, which specify the expression level of gene *i* in cell fate *µ*. The {*ξ*_*µi*_} are an input to our model and can be calculated from single-cell atlases [11–13, 33].

We would like a way to represent the current state of the system not only in gene expression space (*x*_*i*_(*t*)), but also in the space of possible cell fates [33]. Given a set of cell fates {*ξ*_*µi*_}, we can define two different quantities that map the gene expression state *x*_*i*_(*t*) to a *p*-dimensional vector in cell fate space. The first is through defining the overlap, which is equivalent to magnetization in spin systems:

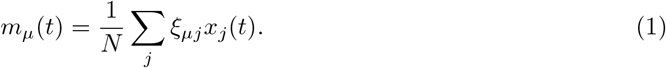

As can be seen from the expression above, the *m*_*µ*_(*t*) are simply the dot products of the current gene expression state with the expression profiles of each of the *p* cell fates we wish to model. While informative, one major drawback of *m*_*µ*_ is that if two cell fates are highly correlated (as is often the case for closely related lineages) the corresponding magnetizations will also be similar.

For this reason, it is useful to also define a second set of quantities we call generalized order parameters *m*^*µ*^ (denoted with an *upper* index) that account for the fact that gene expression profiles of closely related cell fates are correlated [39]. To define this generalized order parameter,first we define the matrix

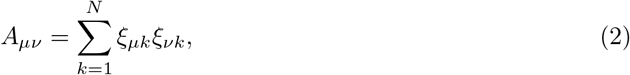

which measures the pairwise similarity between different cell fates. The generalized order parameters can be defined in terms of the inverse of this matrix through the equation

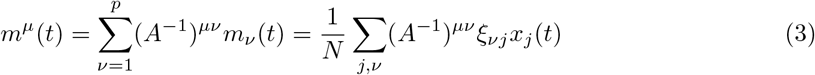

Unlike magnetizations, the generalized order parameters *m*_*µ*_ for two highly related cell fates are generically very different because of the presence of *A*^*™*1^ in the expressions above.

The order parameters *m*^*µ*^ also as a natural coordinate system for cell fate space. This is because Eq. 3 has a geometric interpretation in terms of the linear projection of *x*_*i*_(*t*) onto the *p*-dimensional subspace spanned by the cell type vectors {*ξ*_*µj*_} _*µ*=0,…,*p™*1_ [33]. In the appendix, we show any *x*_*i*_(*t*) can be written as

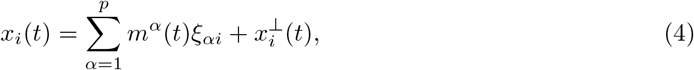

where *m*^*α*^ is the generalized order parameters for cell fate *α* and 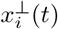 is the part of gene expression vector that is perpendicular to p-dimensional subspace spanned by *{ξ*_*µj*_*}*_*µ*=0,…,*p™*1_ [33]. In other words, the first term represents the gene expression relevant to cell fate, while the second term contains all the gene expression information unrelated to cell type (such as the activity of housekeeping genes). Eq. 4 provides a natural way to transform between the *N*-dimensional gene expression space, *x*_*i*_(*t*), and the *p*-dimensional cell fate space, 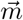 (*t*) = (*m*^1^(*t*), *m*^2^(*t*), …, *m*^*p*^(*t*)) (this corresponds to the top arrows in Figure 1).

To model developmental dynamics, we make use of the analogy between Modern Hopfield networks and developmental landscapes to formulate a dynamic update rule for gene expression of the form

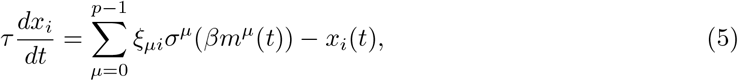

with *τ* a time constant setting the speed of dynamics, *σ*^*µ*^ the soft-max function defined as

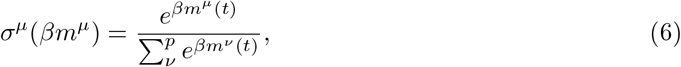

and *β* an inverse temperature parameter that controls the steepness of the non-linearity in the softmax. In the limit of zero temperature, this dynamical update rule ensures that the cell fates {*ξ*_*µj*_} are fixed points. This can be seen by noting that when 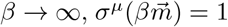 for the direction *µ* = *γ* in cell fate space with the highest magnetization (i.e. 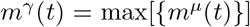) and zero otherwise, 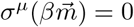 if *µ* ≠ *γ* (see SI section B 2).

We can also rewrite the dynamics in Eq. 5 entirely in cell fate space in terms of the *m*^*µ*^ by noting that

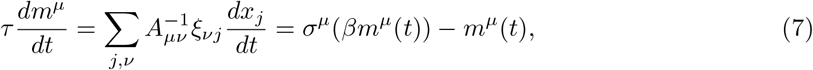

where in the first equality we have used Eq. 3 and in the second equality we have used Eq. 5. This expression shows that our differentiation dynamics do not change gene expression in directions 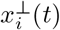 perpendicular to the p-dimensional subspace defined by the cell fates {*ξ*_*µi*_} (i.e.,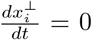). For this reason, we will largely ignore 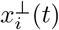 in what follows.

To make a connection with the Waddington landscape and as a prelude to incorporating signaling-induced cell fate transitions, it is helpful to rewrite Eq. 7 in terms of a “potential function” *V* (*t*). In order to do so, we define a potential function of the form

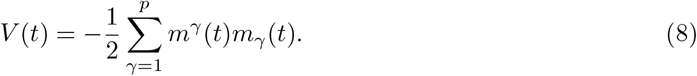

and note that the dynamical Eq. 7 can be rewritten as

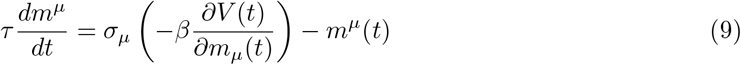

This looks almost like damped gradient dynamics (with the *−m*^*µ*^ acting as a damping, friction-like term), with the key difference that the *p*-dimensional “force” vector 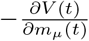 is passed through a non-linear soft-max function.

In cell fate space, the potential *V* is a inverted parabola centered at the origin. In the update rule, the state is pushed by a force caused by this potential, 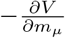, down along this inverted parabola. This potential has no minima, but the dynamics have a steady state because the non-linearity *σ*_*µ*_ bounds *m*^*µ*^ such that |*m*^*µ*^| *≤* 1. The combined effect of the parabola and the bound on *m*^*µ*^ is that a state is pushed down the parabola and then hits the wall represented by the bound, reaching an attractor and remaining there because it can’t be pushed farther along the parabola. This illustrates that we can think of this dynamics as a system pushed by a force defined by the gradient of a potential, but now rectified through a non-linearity.

In the dynamics defined by Eqs 5 and 7, the p-cell fates of interest {*ξ*_*µj*_} are always stable attractors. However, we know that during development cells can differentiate in response to external signals. During a cell fate transition, cells can change their gene expression profiles from an initial attractor, for example a lung progenitor state, to another attractor corresponding to a more differentiated cell fate such as an alveolar cell. To incorporate this in the model, we add a signal-dependent potential 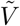 to our potential of the form:

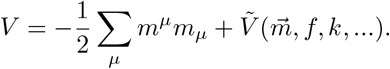

This signal-dependent potential 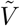 is a function of the cell fate coordinates 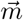 as well as signalling parameters *f, k*, … which represent signals received by the cell. The parameters control the stability of attractors in the landscape. For example, an increase in parameter *f* or *k* might destabilize some attractors and stabilize others. The specific form of 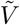 is constructed by considering different classes of decision-making. These classes, described by Sáez et al. [21], Rand et al. [23], Raju and Siggia [24], Camacho-Aguilar et al. [25], are broad categories that capture general features of cell fate transitions, such as the number of attractor states and the possible paths between them. The 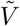 are universal functions constructed from normal forms of bifurcation and hence uniquely identify the decision making class associated with a cell fate transition. Finally, since 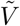 is a scalar function of the vector 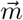, it must be dependent on 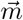 in such a way that every raised index (e.g. *m*^*µ*^) is matched by a lowered index (e.g. *m*_*µ*_) so that the result is a scalar (see section A 3 for a discussion of invariant transforms and the interpretation of these covariant and contravariant vectors). With these additions, the full dynamics of our model are:

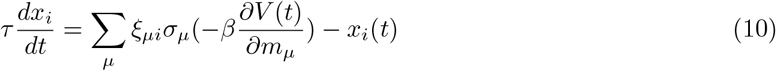

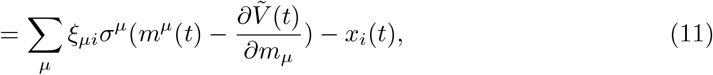

or in cell fate space

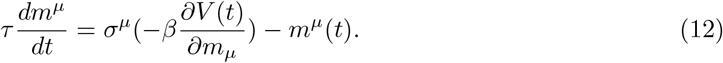

Note, that these dynamics no longer possess a Lyapunov function and the right hand side can no longer be written as a gradient of a potential function due to the soft-max function *σ*^*µ*^.

Our mathematical construction is summarized in Figure 1. The top row of Figure 1 shows the relationship between gene expression and cell fate coordinates. The cell type gene expression profiles *ξ*_*µi*_ provide a way to translate between gene expression *x*_*i*_ and cell fate coordinates *m*^*µ*^. The middle row of figure 1 illustrates how the dynamics between the spaces of gene expression and cell fate are related. As in the dynamics of a Hopfield model, the stored patterns – or cell fates – *ξ*_*µi*_ provide a way to move between the spaces of gene expression and cell fate: 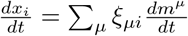. The dynamics 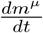 involve the force exerted by the cell fate landscape *V*, which is controlled by the signalling parameters {*f, k*, …}. These parameters define an abstract signaling space, and they represent chemical and mechanical cues that cause the cell to change fates. By featuring these three spaces – gene expression, cell fate, and signaling – our model is able to predict trajectories corresponding to landscapes of any decision-making class.

### B. Constructing potentials from elementary bifurcations

The model described in the previous section combines Hopfield dynamics with a landscape. The question, then, is how to construct a landscape. Landscapes can be defined by the bifurcations they contain. A bifurcation is when a parameter is varied and the stability of a fixed point is changed as a result. The number of parameters one varies to cause the bifurcation is called the codimension of the bifurcation. All local bifurcations of codimension less than or equal to 5 have been enumerated, and they are called elementary catastrophes [17, 40]. Generic forms for landscapes of these elementary catastrophes are called normal forms, and they take the form of polynomials. For example, one of the simplest bifurcations is the codimension-1 fold, or saddle node, bifurcation, which has the normal form *V* (*x*) = *x*^2^ + *a*. *a* is the bifurcation parameter. When *a <* 0, there are two fixed points; when *a >* 0, there are none.

To construct landscapes for cell fate decisions, we follow the method described by Rand et al. [23]. Their work sets the mathematical foundation for identifying universal features of cell fate transitions. We provide a conceptual overview of their method of constructing landscapes. Landscapes are categorized by decision-making classes defined by numbers of attractors, saddle points, and the paths between them. The structure of each class can be depicted as a decision graph showing the paths between attractors and saddle points (see Figure 2). Instead of using all of the elementary catastrophes, these decision-making classes are constructed by combining two simple kinds of bifurcations: folds and heteroclinic flips. These two bifurcations represent the basic ways one can alter decision-making classes: fold bifurcations create or destroy stable fixed points and saddle points, while heteroclinic flips change which stable fixed points are connected. They are sufficient for constructing any class that can be represented in the decision graph structure [23]. Complicated higher-codimension bifurcations like the swallowtail and butterfly catastrophes are neglected because they would require very special circumstances, and hence are assumed to be evolutionary unlikely since they require fine tuning [41].

**FIG. 2:**
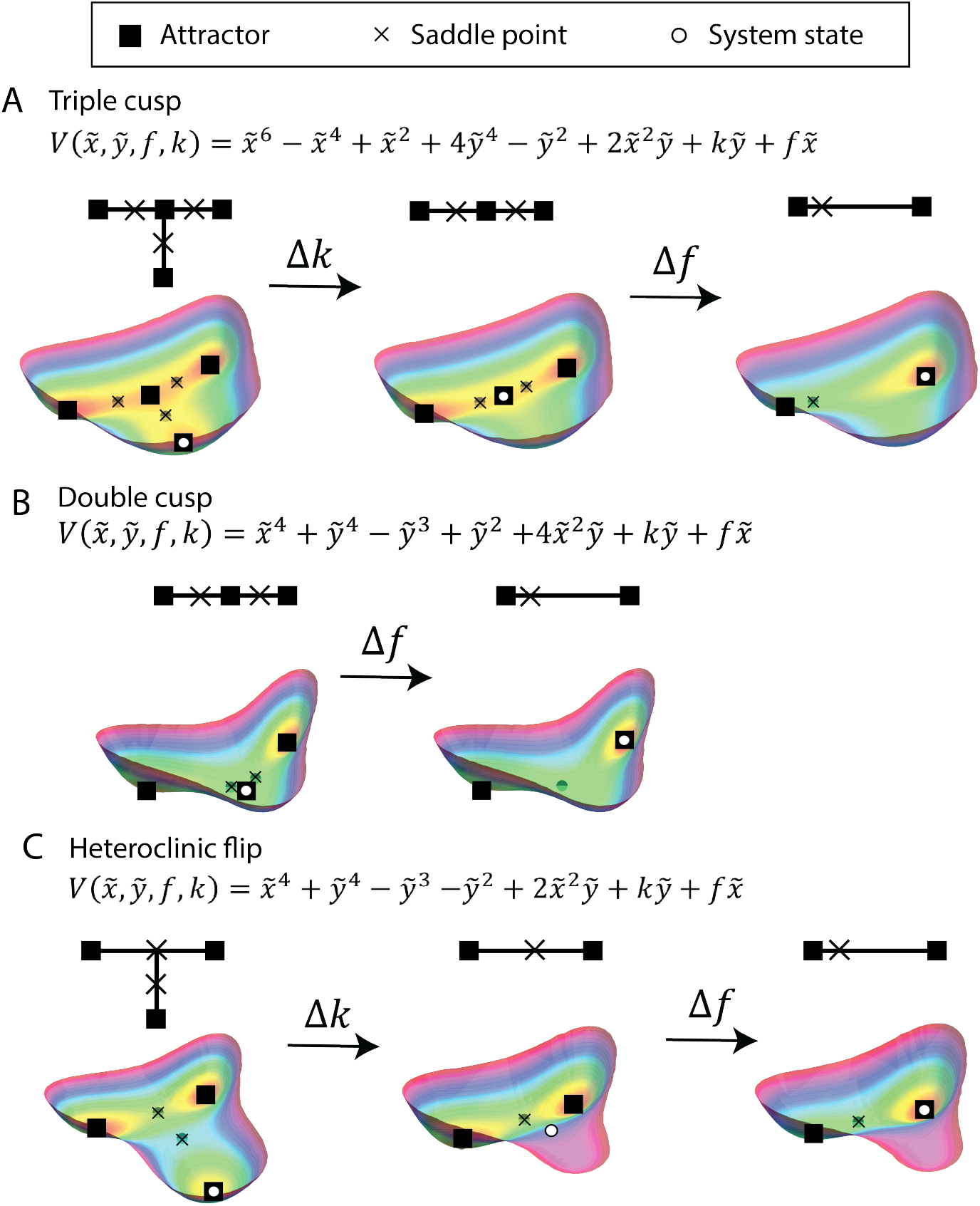
Three cell fate decision-making classes. The potential 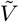 is shown for each two-dimensional land-scape. *f, k* are bifurcation parameters; signaling conditions change these parameters and cause attractors to be stabilized or destabilized. A graph showing the underlying decision structure is shown for each class alongside a representative landscape. A. The triple cusp class. As shown in the decision structure graph, this class contains four attractors and three saddle points. A change in parameter *k* destabilizes the bottom attractor and shifting *f* destabilizes one of the top two attractors. B. The double cusp class consists of three attractors and two saddle points. When *f* is varied, the central attractor is destabilized. C. In the heteroclinic flip, there are three attractors and two saddle nodes. *f* controls the stability of the bottom attractor while *k* tilts the landscape towards one of the top attractors.

Formally, this argument relies on the properties of gradient-like Morse-Smale systems. These are dynamical systems that are well-behaved: they are structurally stable and every trajectory ends in a fixed point [24]. These properties match the picture of the Waddington landscape, where every cell attains a cell fate in a robust, controlled manner. Structural stability means that trajectories don’t change drastically with small perturbations, but in this case this only applies to the parameters not involved in bifurcations. Small changes in the bifurcation parameters can cause significant changes in the system’s behavior. A bifurcation causes a transition from one Morse-Smale system to another. It has been shown that the only bifurcations needed to transition between similar Morse-Smale systems are the fold and heteroclinic flip [24, 42].

The equations describing the landscapes for these classes are based on the normal forms of the elementary catastrophes. In other words, one takes the unfolding of a sufficiently high dimension and adjusts the parameters of terms to match the behavior of the decision graph. For example, the elliptic umbilic 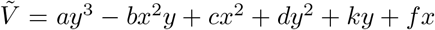 serves as the starting point for writing out the landscapes described below (the corresponding equations are shown in Figure 2) [23]. The landscapes below are found by modifying this normal form in two ways. The first modification is to add higher-order even terms (*x*^4^, *y*^4^, *x*^6^, *y*^6^) so that 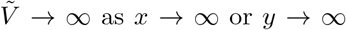. This puts the system in a potential well so that it doesn’t escape to infinity. The second modification is to choose the coefficients of each higher-order term (*a, b, c, d*) to match the behavior of the chosen decision graph. After this adjustment, the only bifurcation parameters are the coefficients of the linear terms *f, k*. These represent the two parameters controlled by signals received by the cell. Their effect mathematically is to tilt the landscape along each of the axes. As shown in Figure 2B for the case of the triple cusp landscape, tilting along the *y*-axis corresponds to a change in *k* that destabilizes the initial attractor basin. A change in *f* tilts the landscapes along the *x*-axis which biases the decision towards one of the final two fates. It is possible to define these parameters explicitly: for example, one could define each one as a function of the amount of Notch ligand produced by a cell’s neighbors and received by a cell’s receptors. However, for the purposes of this paper, we do not explicitly define *f, k*. For the sake of generality, we define them as abstract parameters that determine the stability of attractors. In simulations, we control these variables manually, as described in the Supplementary Information section B 1.

We now describe some specific cases of decision-making classes. These are characterized by their corresponding decision graphs, which show the attractors, saddle points, and paths between them. Classes with one or two attractors are trivial. Interesting decision-making dynamics occur with three or more attractors because this represents a cell making a decision between two fates as it leaves its initial state. The two simplest classes with three attractors are the double cusp and the heteroclinic flip.

The double cusp is shown in Figure 2A. It consists of three attractors and two saddle points, with saddle points separating the initial attractor from each of the final two. A cusp bifurcation is like two fold bifurcations which intersect at the saddle point. One can build a double cusp using four intersecting fold bifurcations. Shifting the bifurcation parameter *f* destabilizes the initial attractor basin and biases the cell towards one of the final two fates. The heteroclinic flip is show in Figure 2C. It also has three attractors and two saddle points, but the saddle points are arranged such that they are connected to one another. With this arrangement, the parameter *k* controls the stability of the initial attractor independently of *f*, which biases the cell towards one of the two final fates.

A third, more complicated class is shown in Figure 2B. This class includes a fourth attractor in the middle of the three primary attractors. This corresponds to a transient attractor which is a mixed state of the final two attractors. When *f* is varied, the cell moves from the initial attractor to this intermediate attractor, and *k* destabilizes this intermediate state to push the cell to one of the final two states. We include this class because there is evidence to suggest that some cell fates have a bipotent progenitor state which is a mixture of the final fates [33, 43].

### C. Identifying experimental signatures of decision-making classes

Once we have constructed a landscape for a given class, we can use our Hopfield-like dynamics to simulate cell fate transitions corresponding to that class. Figure 3 shows the trajectories of these simulations in cell fate space. By comparing these trajectories across landscapes, we can identify signatures for each decision-making class. In the double cusp, signals push the cells directly to their final fates so the trajectories for this class are straight lines. In the triple cusp, cell spend more time in the intermediate state than in the other states along the trajectory because it is transiently stable. As a result, there is a higher density of cells in a localized patch on this trajectory. This intermediate cluster is the defining feature of the triple cusp class. Finally, the heteroclinic flip can be identified by its curved paths, which cells take as they move away from the second saddle node. Although the details of the landscape can vary the appearance of these features, they can be used to distinguish between classes. By searching for these signatures in actual data, it’s possible to find signs of decision-making classes and directly compare real transitions to theoretical landscapes.

**FIG. 3:**
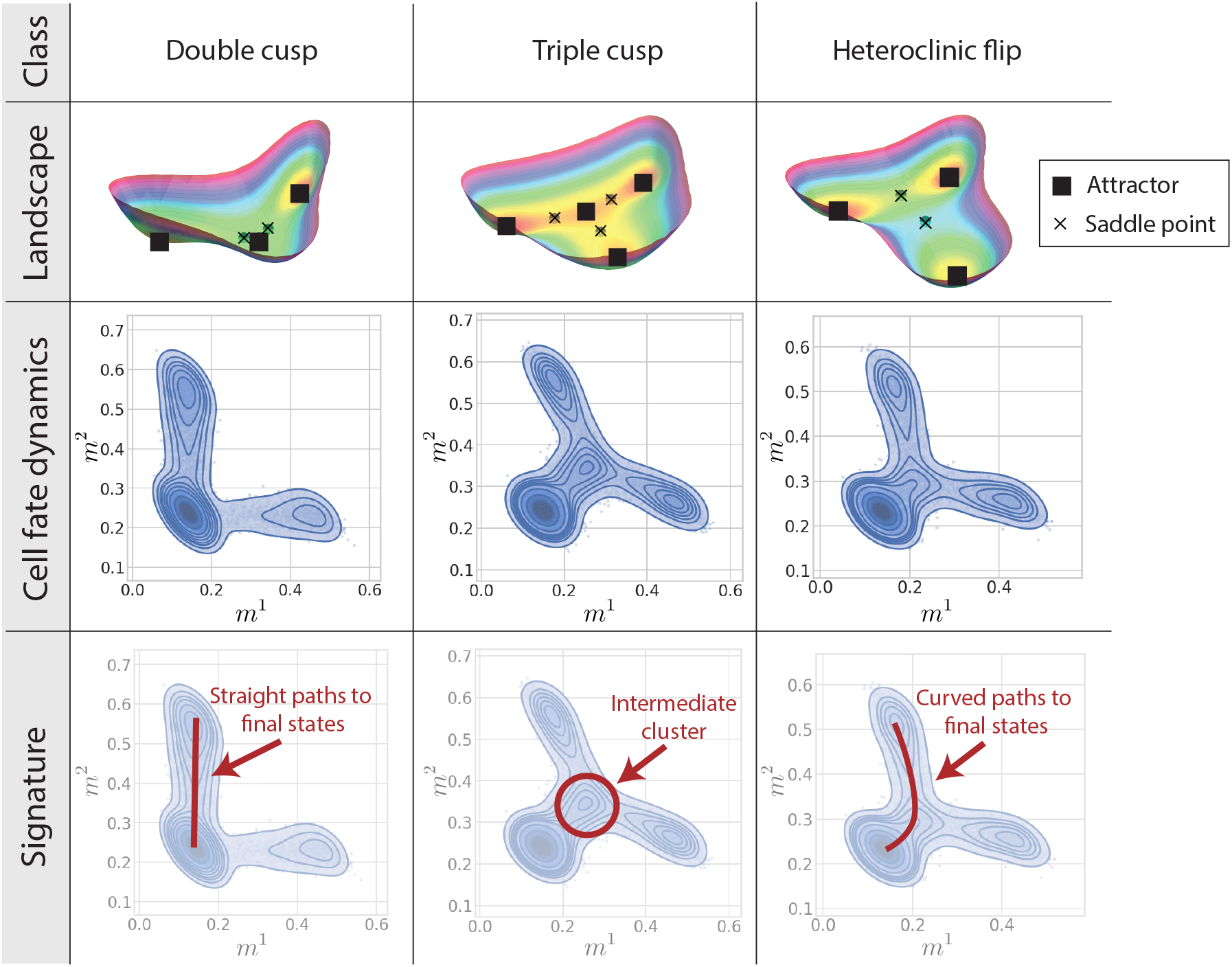
Three decision-making classes and their signatures in cell fate space. For each class, and example landscape is shown, with attractors marked by boxes and saddle points marked by “x”s. Contour plots of trajectories for each class are shown, plotted on cell fate coordinates corresponding to the two final fates (see Figure 2 for scatter plots of similar trajectories). The bottom row highlights the distinctive features of each class. The double cusp contains straight lines from the first cell type to the final two types. The triple cusp contains a distinct, intermediate cluster corresponding to cells spending time in the transient attractor. The heteroclinic flip features paths that curve around the saddle points, leading to the final states.

It is possible to measure the cell fate coordinates *m*^*µ*^ in real systems by applying scTOP to scRNA-seq data [33]. Time-series data is particularly valuable for observing the trajectory of cells, as identifying some features of dynamical systems requires temporal information. For example, knowing how much time cells spend in different states allows one to distinguish between transient and stable attractors. In the case of embryonic development, snapshots across developmental time points show the path of cells as signaling and mechanical conditions are varied. In this section, we look for signatures of decision-making classes in two contexts: *in vitro* mouse hematopoiesis and the developing mouse lung.

#### 1. Cell fate decisions in hematopoiesis

Blood requires frequent replenishment, so a wide range of blood cells are regularly created in bone marrow. These blood cells originate from hematopoietic stem and progenitor cells (HSPCs), which have the capacity to differentiate into many blood cell fates. They are responsible for maintaining stable populations of blood cells. While many different cell fate hierarchies have been proposed for hematopoiesis, lineage tracing data from Weinreb et al. [43] suggests that HSPCs primed to become different cell types may exist on a continuum rather than passing through distinct progenitors for each mature fate. For example, some HSPCs which become monocytes take paths which are closer to dendritic cells while others take neutrophil-like paths. By comparing populations of HSPCs and the paths they take in cell fate space, our model can identify signs of landscapes associated with these possible scenarios.

Lineage tracing experiments in Weinreb et al. [43] enable a precise analysis of progenitors and their eventual fates. HSPCs were extracted from the bone marrow of mice and tagged with DNA barcodes unique to each cell. These barcodes are retained when cells divide. Since scRNA-seq destroys the cell it samples, it’s difficult to observe the gene expression of a particular cell over time. Lineage tracing by barcoding is one way to bypass this difficulty, since it allows mature cells to be matched to clonal progenitors via the barcode. In this experiment, after cells were barcoded and allowed to proliferate ex vivo, some cells were sampled, and some were replated and allowed to divide and differentiate further. This process was repeated twice, which led to samples being taken 2, 4, and 6 days after barcoding (see Figure 4 A).

**FIG. 4:**
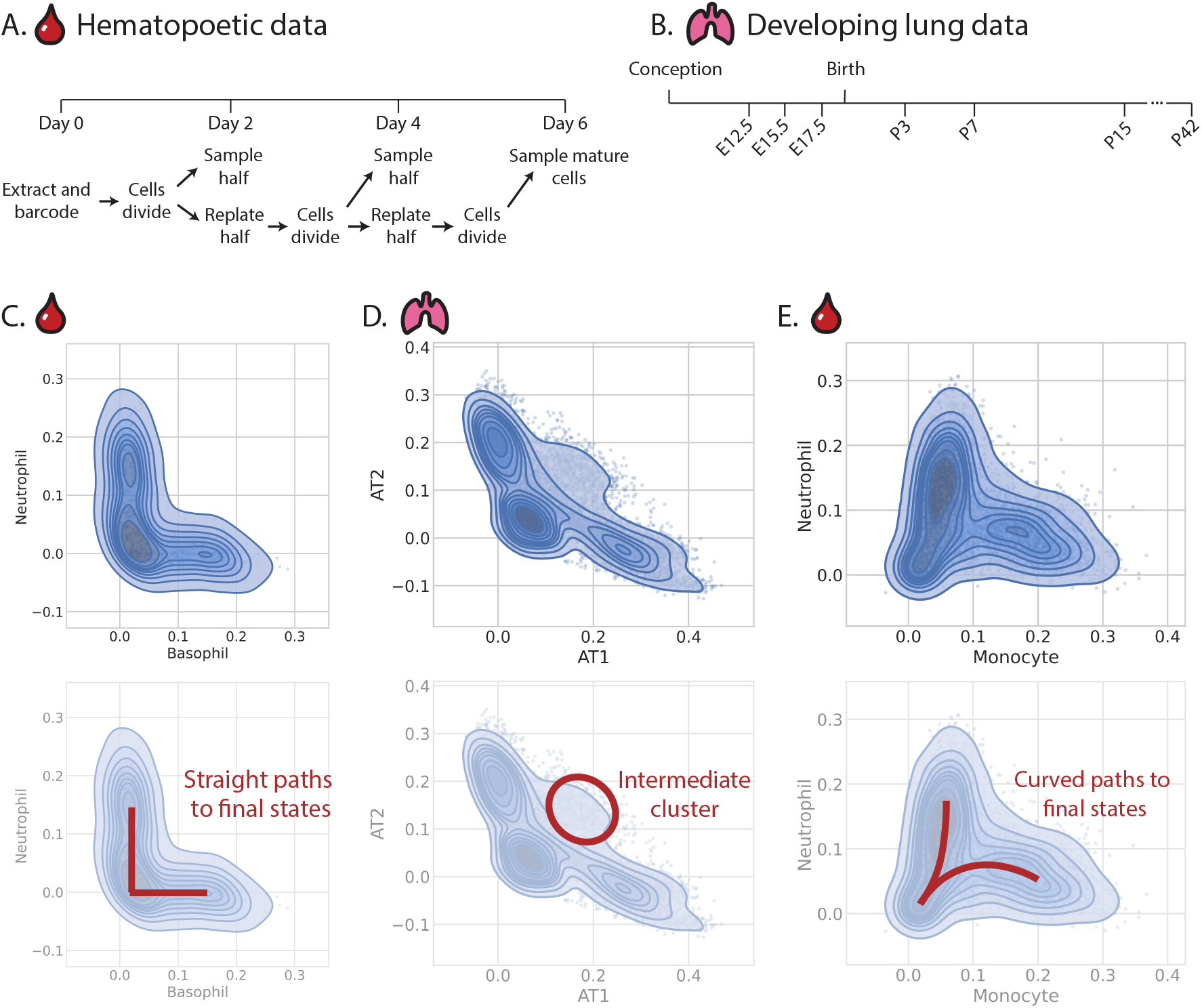
Experimental scRNA-seq data and corresponding signatures of landscapes. A. In Weinreb et al. [43], hematopoietic progenitor cells were extracted from the bone marrow of mice and tagged with DNA barcodes. Then they were allowed to divide ex vivo. On days 2 and 4 after extraction, some cells were sampled with scRNA-seq while others were allowed to proliferate. On day 6, mature blood cells were sampled. B. Zepp et al. [46] collected lung samples across stages in mouse lung development, from embryonic to adult mice. The samples begin at embryonic day 12.5 (E12.5, i.e. 12.5 days post-conception) and end at post-natal day 42 (P42). (C,E) The hematopoietic data is plotted on cell fate axes. Only cells belonging to clonal families of the relevant bipotent progenitors are shown in each figure. C. Basophil-neutrophil progenitors differentiating, plotted on cell fate coordinates corresponding to basophil and neutrophil alignment. The trajectories are straight, suggesting the signature of a double cusp class. D. The developing lung data is shown in cell fate space. The axes corresponding to AT1 and AT2 cell fate coordinates. The presence of an intermediate cluster suggests a triple cusp class. E. Monocyte-neutrophil progenitors maturing, shown on axes corresponding to monocyte and neutrophil cell fate coordinates. The cells take curved paths to their eventual fates, suggesting the presence of a saddle point and therefore a heteroclinic flip class. The corresponding scatter plots, colored by time point, are shown in Figure 7.

HSPCs have many possible fates, and landscapes that involve many attractors can be highly complex. To compare this data with landscapes containing only three attractors, we limited our analysis to bipotent progenitors. In other words, only clonal families which involved exactly two final fates were included. Figure 4C, E shows basophil-neutrophil and monocyte-neutrophil bipotent populations. The basophil-neutrophil cells take straight paths to their eventual fates, which suggests a double cusp class. In the case of differentiating monocyte-neutrophil progenitors, the trajectories are curved like cells in a heteroclinic flip class approaching and then diverging away from a saddle point. The neutrophil-like path for monocytes described in Weinreb et al. [43] could be the result of a saddle point between monocytes and neutrophils.

Identifying signatures of these classes in hematopoiesis can aid in verifying the lineage tree of differentiation. If two fates exhibit a dual-cusp-like path, they are more distantly-related; in fact, it’s possible to take two entirely unrelated cell fates and observe the dual cusp, since there will be no correlation between them. On the other hand, classes like the heteroclinic flip and triple cusp indicate closer relations, which may indicate that the cell fates are adjacent on the lineage tree.

#### 2. Cell fate decisions in the developing lung

Displaying a variety of specialized cell types, the lung provides a rich system for studying cell fate decisions. Different transitions between these types are possible in homeostasis, injury, and transplant [44]. In murine embryonic development, cells in the foregut specify into lung epithelium around 9 days post-coitum (dpc) [45]. The alveoli, where gas is exchanged between the organism and the environment, appear around 16.5-18.5 dpc. Eventually these cells become alveolar type 1 (AT1) and alveolar type 2 (AT2) cells. The lung continues to remodel and develop even after birth, as the newborn animal begins to breath air [46].

Zepp et al. [46] performed scRNA-seq on samples of mouse lungs from 12.5 dpc to 42 days after birth to investigate this complex process (see Figure 4 B). To identify the landscape involved in alveolar specification, we analyzed all samples in this dataset labelled by the authors as alveolar progenitors, AT1 cells, and AT2 cells. For these cells across all time points, the cell fate trajectories for alveolar cells are shown in Figure 4 D. Many cells follow curved paths suggestive of a heteroclinic flip, but the population of cells in a mixed AT1/AT2 state indicates the signature of a triple cusp. The triple cusp can behave like the heteroclinic flip in some regions of signaling space. In other words, depending on the timing of signals received by the cell, it may be induced to follow one transitional path or another. Although it is possible to achieve the heteroclinic flip trajectory with a landscape containing triple cusps, it is not possible to do the reverse. Thus, it is more likely that the alveolar differentiation resembles the triple cusp.

By identifying a signature of the triple cusp class, we learn about how the AT1 and AT2 fates are related and the possible existence of a temporary mixed state. Additionally, each cell experiences a different landscape according to its external environment. In the population that passes through the transient attractor state, the cells move toward mature AT1 and AT2 fates shortly after birth. This may indicate that the remodeling associated with air-breathing destabilizes the intermediate attractor seen in the triple cusp.

## III. DISCUSSION

The theory of dynamic systems captures the phenomenology of cell fate transitions with minimal parameters. Even if the true epigenetic landscape is highly complex, we can use generic decision-making classes to describe the local nature of the landscape. Instead of requiring exact information of how each gene affects every other gene, the geometric picture shifts the focus to the most general features of transitions: attractors, saddle points, and flows. This abstraction allows us to identify classes of cell fate transitions and make progress towards finding general principles underlying these transitions. Our landscape-driven model preserves the benefits of abstracting to a low-dimensional cell fate space while also providing a way to dialogue with scRNA-seq experiments. Our model avoids the need for precise knowledge of gene network interactions by leveraging the pattern-retrieval capability of Hopfield networks. However, it is worth emphasizing the ambiguities inherent in this formulation: decision-making classes do not fully constrain the details of dynamical trajectories (see section C for further discussion).

By combining bifurcation theory and modern Hopfield networks, our model connects the spaces of gene expression, cell fate, and signaling. The order parameters *m*^*µ*^ from Yampolskaya et al. [33] provide common axes on which to compare simulations and real experiments, enabling direct comparison between theory and data. Simulations for landscapes containing triple cusp, double cusp, and heteroclinic flip bifurcations revealed trajectory signatures for identifying these bifurcations. By analyzing time series scRNA-seq data in cell fate dynamics, we identified signatures of classes in alveolar development and hematopoietic differentiation. This approach found evidence for a double cusp in basophil-neutrophil differentiation, a heteroclinic flip in neutrophil-monocyte differentiation, and a triple cusp in alveolar differentiation.

With these methods of analysis alongside the abundance of time-series scRNA-seq atlases, it is now possible to group cell fate transitions across biological contexts according to decision-making class. Classifying cell fate decisions can aid in determining how fates are related and which transitions are possible. With enough information, our model may be able to predict the signals necessary to take the path from one identity to another.

With our model, it is possible to explain a wide range of experimental observations which we do not cover in this introductory paper. For example, more work can be done in developing meaningful signaling parameters. In the alveolar development example shown in figure A 1A, the parameters that shifted the landscape represented the sum of chemical and mechanical effects when the alveoli formed and the mouse began breathing after birth. Future *in vitro* work could explore the cell fate trajectories of cells in response to signals by extracting endogenous cells, applying signaling factors, and taking scRNA-seq data at time points throughout the process. This would allow for more accurate quantitative landscape-based modeling of these transitions by directly relating the effects of signals to displacements in cell fate space. The application of either chemical gradients or mechanical forces to cell populations that are repeatedly sampled would allow our models to predict the effects of these changes on measurable axes.

## IV. METHODS

### A. Defining cell fate coordinates

As previously described in Yampolskaya et al. [33], generalized Hopfield order parameters (called single-cell Type Order Parameters (scTOP)) can be applied to gene expression data to define cell fate space. In the case of a single cell, let *x*_*i*_ denote the cell’s expression of gene *i*. These are continuous values because, in scTOP, the RNA counts are converted to z-scores reflecting the rank-ordering of genes within a cell (e.g a gene *i* at the 50 percentile is assigned *x*_*i*_ = 0, a gene j at 84.6 percentile an expression level of *z* = 1, etc). Let *ξ*_*µi*_ be the expression of gene *i* in cell type *µ*. With scTOP, *ξ* is a list of gene expression profiles of known cell types. These are derived from scRNA-seq atlases such as the Mouse Cell Atlas and Tabula Sapiens by averaging across populations of cells, where each population corresponds to a different cell type [11, 12]. Then, as introduced by Kanter and Sompolinsky [39], the generalized Hopfield order parameters are defined as follows:

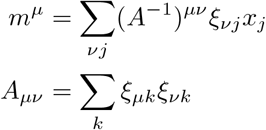

where *m*^*µ*^ is the order parameter which measures alignment with the *µ*th cell type, and *A* is the matrix of correlations between each cell type.

The order parameters *m*^*µ*^ have a natural interpretation as a set of coordinates because they are equivalent to a projection onto the non-orthogonal subspace spanned by known cell types. The cell types *ξ* form a subspace of gene expression space. When a cell’s RNA counts are measured through scRNA-seq, this represents another vector in gene expression space. By projecting this vector onto the cell type subspace, we can describe it with a new set of coordinates, where each axis measures alignment with a cell type. In other words, 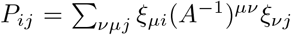 is a projection matrix that transforms vectors from gene expression space (denoted by *j*) to cell type space (denoted by *µ*). We can rewrite the original vector describing the gene expression of the cell as a decomposition onto this subspace:

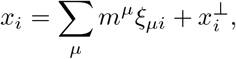

With

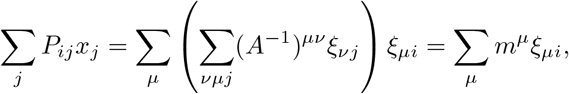

And

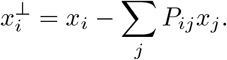

In the equation above, one can see that *m*^*µ*^ are the coefficients of decomposition onto the *ξ* vectors, and 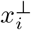 is the component of *x*_*i*_ that is perpendicular to the cell type subspace. This perpendicular component corresponds to gene expression information about processes that are not indicative of cell type, such as the expression of housekeeping genes. It corresponds to the information that is lost when going from the higher-dimensional space of gene expression to the lower-dimensional space of cell fate. In other words, 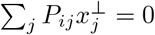. This follows from the definition above by noting that for a projector *P* ^2^ = *P*. In the classic Hopfield network, the stored patterns *ξ*_*µi*_ are orthogonal, in which case 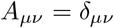 and 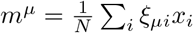

Since these generalized order parameters make use of a change of basis (from gene expression space to the basis of cell fate vectors), we use Einstein summation notation. The covariant and contravariant forms of the order parameters are distinguished by upper and lower indices. In this notation, we have a contravariant vector *m*^*µ*^, a covariant vector *m*_*µ*_, and a metric tensor *g*_*µν*_ that is dependent on the correlations between patterns.

We define the metric tensor using the product of our new basis vectors, *g*_*µν*_ = *A*_*µν*_ = *ξ*_*µ*_ *ξ*_*ν*_, with the inverse of the metric tensor defined with upper indices: 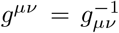. For the covariant form (with a lowered index), we define 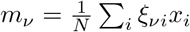. For the contravariant form (with a raised index), we define:

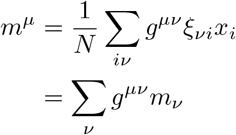

### B. Choice of landscapes

With this formulation, it is possible to simulate dynamics associated with any cell fate landscape. We consider landscapes associated with decision-making classes containing three primary attractors: a progenitor state with two possible mature fates. For simplicity and ease of visualization, this paper considers two-dimensional landscapes, 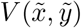. However, in this model, three primary attractors corresponds to a three-dimensional cell fate space, since each dimension is associated with a cell fate. To align the attractors of a two-dimensional landscape with three-dimensional cell fate space, it is necessary to rotate the plane of the potential to align with the attractors. The details of transforming from 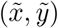 space to (*m*^0^, *m*^1^, *m*^2^) space are explained in A 2.

The gradient of the landscape 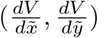 is a function of the signaling parameters (*k, f*), which control the bifurcations. The values for the signaling parameters (*k, f*) were varied as follows. For the landscape with a triple cusp, the system starts with *k* = 0, *f* = 0. The signaling parameter *k* is increased from 0 to 0.3, destabilizing the attractor in which the system started. Then, *f* is changed from 0 to 0.3, causing the intermediate attractor to disappear and tilting the landscape towards one of the final fates. In the landscape with a double cusp bifurcation, the parameters start at *k* = 0.15, *f* = 0. Parameter *k* is kept at 0.15, and *f* is changed from 0 to 0.1. This destabilizes the initial attractor and pushes the cell towards one of the final cell fates. The landscape with a heteroclinic flip bifurcation starts with *k* = 0.5, *f* = 0. Signal *k* is shifted from 0.5 to 2 to destabilize the initial attractor, and before that bifurcation is fully complete, *f* is changed from 0 to 0.5 to tilt the landscape towards the cell fate on the right-hand side. A 2.

### C. Simulation Details

To simulate Figure 3, we used scRNA-seq samples of alveolar type 1 and type 2 cells from Herriges et al. [47] and early epithelial samples from Negretti et al. [48] to define the three cell fates (two mature fates and one progenitor state) for each of the three landscapes.

The initial and final values for the signaling parameters (*k, f*) are specified for each simulation, along with the start and stop times for each signal. The signaling parameters are gradually changed from the initial to final values over the course of this defined signaling window (see SI section B 1 for more details). It uses this information, along with the attractor states *ξ*_*µi*_, to calculate the update step:

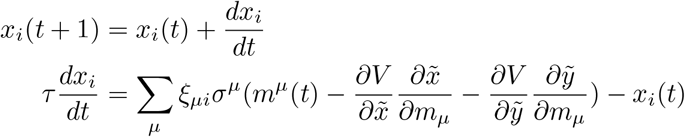

The derivatives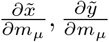 come from applying the chain rule, and their specific forms are derived in SI section A 2. For each choice of landscape *V*, the simulated cell begins in the progenitor cell fate. Then, the signaling parameters *f, k* are changed to destabilize the initial attractor state. Due to this bifurcation, the cell is pushed from its initial state. The parameter *f* tilts the landscape towards one of the terminal fates, and the cell ends in the corresponding attractor basin. Please see Supporting Information for details.

## V. DATA AVAILABILITY

scTOP is available as a package on the Python Package Index (https://pypi.org/project/scTOP/) and the code is accessible on Github (https://github.com/Emergent-Behaviors-in-Biology/scTOP). The scRNA-seq data used in this paper come from a variety of sources. The alveolar lung data used for simulations is available on NCBI’s Gene Expression Omnibus under accession code GSM6046035, as well as the Kotton Lab’s Bioinformatics Portal (http://www.kottonlab.com) [47]. The early lung epithelial sample used as the simulated progenitor state is available under accession code GSE165063 [48]. The embryonic mouse lung data taken from E12.5 to P42 is available under accession code GSE149563 [46]. The hematopoietic lineage tracing data is available under accession code GSE140802, and the metadata can be found on Github (https://github.com/AllonKleinLab/paper-data/tree/master/Lineage tracing on transcriptional landscapes links state to fate during differentiation) [43].

## VI. CODE AVAILABILITY

The Python and Mathematica notebooks used for the simulations and figures in this paper are available on Github (https://github.com/Emergent-Behaviors-in-Biology/Hopfield-landscapes).

## Acknowledgments

We acknowledge useful discussions with Jason Rocks, Michael Herriges, and members of the Kotton Lab and Mehta group. The work was funded by grants from the Boston University Kilachand Multicellular Design Program, Chan-Zuckerberg Investigator grant to PM, and NIH NIGMS 1R35GM119461 to PM.

