## Supplemental Information for "Finding signatures of low-dimensional geometric landscapes in high-dimensional cell fate transitions"

#### Contents

**Supplemental Information** 1
A. Mathematical details 1
1. Choice of potential 1
2. Landscape coordinate transformation 3
3. Correlated cell types and invariant transformations 5
4. Potentials over time 7
B. Simulation details 7
1. Signaling dynamics 8
2. Inverse temperature 10
3. Noise 10
4. Robustness to update schedule 10
C. Limitations of the model 12

**References** 12

### A. Mathematical details

#### *1. Choice of potential*

In choosing possible landscapes, we only considered landscapes containing three attractors. As explained in Rand et al. [1], two of the simplest bifurcations with three attractors are the binary cusp and the heteroclinic flip. However, in the case of AT1/AT2 differentiation, we noticed that there appeared to be a transient attractor. To account for this bifurcation, we also included the triple cusp. Bifurcation diagrams for these three bifurcations are shown in Figure 2.

Although these three types of bifurcations can occur in high dimensions, locally they are equivalent to two-dimensional normal forms; in other words, local models for bifurcations are universal, so we expect a two-dimensional landscape to capture the important qualities of a binary decision [1]. For the binary cusp and heteroclinic flip, we use the same equations as Sáez et al. [2]. We found the equation for the triple cusp by introducing a higher-order term, $\tilde{x}^6$, and modifying the coefficients of the lower-order terms to produce behavior matching the decision graph. The equations for all three landscapes are shown in Fig. 2. These landscapes are functions of two variables, $\tilde{x}, \tilde{y}$, and the way we translate between these variables and cell fate coordinates is described in Section A 2

Small changes to the coefficients of these forms modifies the positions of attractors and saddle nodes, which can change the shape of the landscape. For example, changing the coefficient of the $\tilde{x}^2\tilde{y}$ term in the double cusp shifts the positions of the saddle nodes, as shown in Figure A 1. Varying $c$ slightly in this way doesn't change the bifurcation, but it will change the path a ball would take in this landscape and the necessary push needed for the ball to escape the first basin. In terms of cell fate, this corresponds to the strength of signal needed to transition from the initial fate to one of the final fates. Additionally, one can imagine an asymmetry between fates, where less signal is needed to transition to one of the final fates relative to the other. The class of bifurcation does not constrain the possible relative positions of attractor points and saddle nodes.

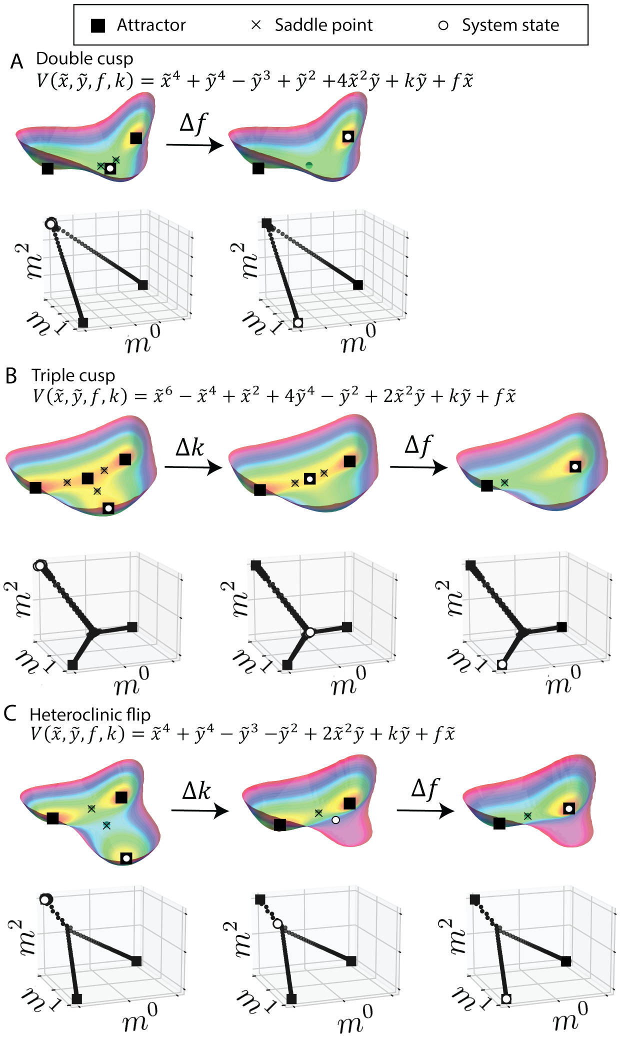


FIG. 1: Simulations of cell fate dynamics for three possible landscapes as they undergo bifurcations. Each landscape is colored by depth to emphasize attractor basins. Below each landscape, the full trajectory in cell fate space ($m^0, m^1, m^2$ for cell fates 1, 2, 3) is shown in black, with white circles indicating the cell states corresponding to the white circles in the landscapes above. The equations for each landscape are shown in terms of variables $\tilde{x}, \tilde{y}$ and signaling parameters $f, k$ (see SI section A 2 for explanation of these coordinates and how they relate to $m^0, m^1, m^2$). For the bifurcation diagrams for each case, see figure 2.

Large changes to the coefficients results in different bifurcation diagrams. As shown in Figure 2, there are regions of parameter space where all three landscapes have the same bifurcation diagrams.

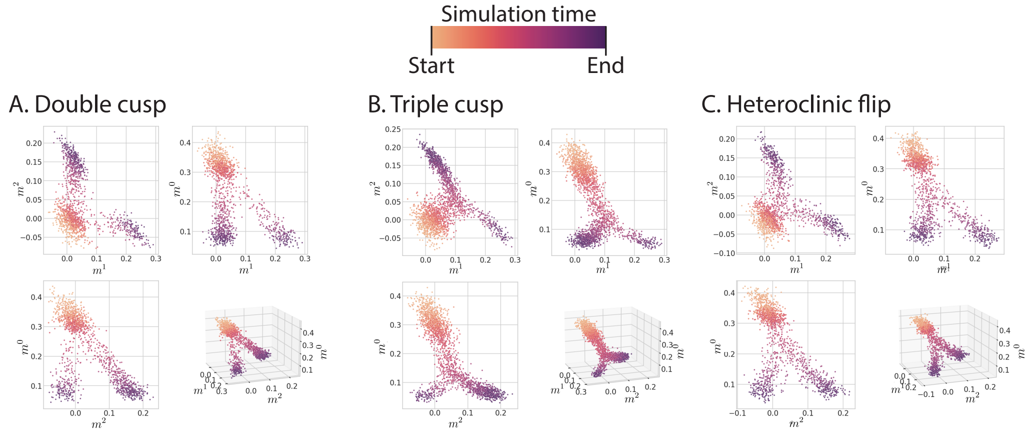


FIG. 2: Simulations of cell fate dynamics corresponding to each of the three classes. The simulations are run similarly to Figure 3, except noise has been added to imitate measurement noise in scRNA-seq data.

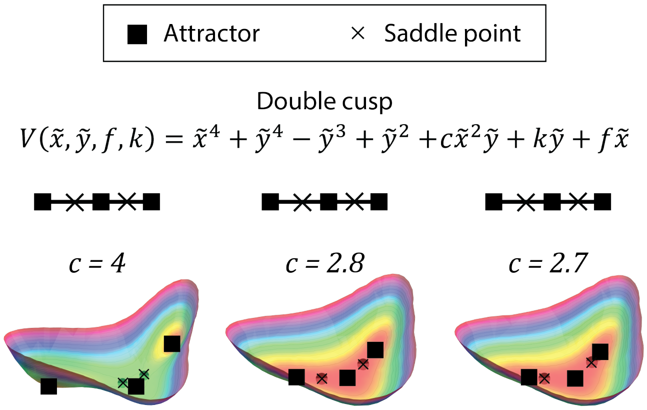


FIG. 3: In the normal form of the double cusp, changing parameter $c$ varies whether the saddle nodes are closer to the initial attractor basin or the final basins. Note that the bifurcation diagram stays the same, since the diagram only conveys a topological description of the bifurcation and not the details of position.

#### 2. *Landscape coordinate transformation*

The normal forms corresponding to each decision-making classes take the form of a 2-dimensional potential, $V(\tilde{x},\tilde{y})$. In our model, alignment with each cell type is given its own axis, set by the Hopfield-inspired order parameters $m^{\mu}$. To align the attractors of the landscape $V(\tilde{x},\tilde{y})$ with the attractors of the Hopfield model, we must transform coordinates from landscape space $\tilde{x}, \tilde{y}$ to cell type space $m^\mu$ (see Figure A 2).

In the case of three attractors (points labelled $a_0, a_1$, and $a_2$), we want to transform from the two dimensions $\tilde{x}, \tilde{y}$ to three dimensions $m^0, m^1, m^2$. For the normal forms of the three bifurcations considered in this paper, the coordinates of these attractors in landscape space $(\tilde{x}, \tilde{y})$ adhere to the following format:

$$\begin{aligned}\tilde{a}_0 &= (0, \tilde{y}_0)\\ \tilde{a}_1 &= (\tilde{x}_1, \tilde{y}_1)\\ \tilde{a}_2 &= (-\tilde{x}_1, \tilde{y}_1)\end{aligned}$$

In other words, one attractor is on the $\tilde{y}$-axis and the other two are symmetrically opposed across the $\tilde{y}$-axis. In cell type space $(m^2, m^1, m^0)$, we want the attractors to be on the edges of the 3-simplex:

$$\begin{aligned}a_0^m &: (0, 0, 1)\\ a_1^m &: (0, 1, 0)\\ a_2^m &: (1, 0, 0)\end{aligned}$$

To facilitate writing this transformation in matrix format, we will move to four-dimensions such that $(\tilde{x}, \tilde{y})$ coordinates become $(\tilde{x}, \tilde{y}, 0, 1)$ and $(m^2, m^1, m^0)$ coordinates become $(m^2, m^1, m^0, 1)$. The index for summation over this four-dimensional space will be given by $\lambda$. Note that once the transformation is applied, it is simple to move to three-dimensional space ($\lambda = 0, 1, 2, 3 \to \mu = 0, 1, 2$) by simply truncating the fourth dimension (for equations involving this transform, if $\mu$ is used for summation it implies the 3-dimensional vector is extended to 4 dimensions, with a value of 1 in the fourth dimension and a subsequent summation over $\lambda = 0, 1, 2, 3$).

The 4-by-4 matrix transforming from $(\tilde{x}, \tilde{y}, 0, 1)$ to $(m^2, m^1, m^0, 1)$ is given by $K$. To compose $K$ such that $\vec{a^m} = K\vec{\tilde{a}}$, the following transformation steps are necessary:

1. Scaling with matrix $S$ such that the triangle formed by $a_0, a_1, a_2$ becomes equilateral with side lengths $\sqrt{2}$. $S = \begin{pmatrix} \frac{\sqrt{2}}{2x_1} & 0 & 0 & 0\\ 0 & \frac{\sqrt{3/2}}{y_0 - y_1} & 0 & 0\\ 0 & 0 & 1 & 0\\ 0 & 0 & 0 & 1\end{pmatrix}$
2. Translating the $a_0, a_1, a_2$ triangle with matrix $T_1$ such that the midpoint of the bottom side is a distance $\frac{\sqrt{2}}{2}$ away from the origin. $T1 = \begin{pmatrix} 1 & 0 & 0 & 0\\ 0 & 1 & 0 & -\frac{\sqrt{2}}{2} - \frac{y_1\sqrt{3/2}}{y_0 - y_1}\\ 0 & 0 & 1 & 0\\ 0 & 0 & 0 & 1\end{pmatrix}$
3. Rotating about the third axis by an angle of $\frac{3\pi}{4}$ with matrix

$$R_1 = \begin{pmatrix} \cos\left(\frac{3\pi}{4}\right) & -\sin\left(\frac{3\pi}{4}\right) & 0 & 0\\ \sin\left(\frac{3\pi}{4}\right) & \cos\left(\frac{3\pi}{4}\right) & 0 & 0\\ 0 & 0 & 1 & 0\\ 0 & 0 & 0 & 1\end{pmatrix}$$

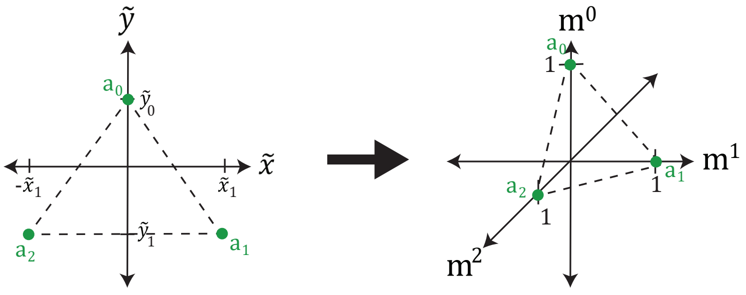


FIG. 4: Transforming from normal form coordinates $\tilde{x}, \tilde{y}$ to cell type coordinates $m^0, m^1, m^2$ to align landscape attractors with Hopfield attractors

4. Rotating by an angle of $\arcsin\left(\sqrt{\frac{2}{3}}\right)$ about the line that crosses through (1, 0, 0, 1) and (0, 1, 0, 1) with matrix $M = T^{-1}X^{-1}Y^{-1}ZYXT$. $T, X, Y, Z$ are defined as follows:

$$T = \begin{pmatrix} 1 & 0 & 0 & -1 \\ 0 & 1 & 0 & 0 \\ 0 & 0 & 1 & 0 \\ 0 & 0 & 0 & 1 \end{pmatrix}$$

$$X = \begin{pmatrix} 1 & 0 & 0 & 0 \\ 0 & 0 & -1 & 0 \\ 0 & 1 & 0 & 0 \\ 0 & 0 & 0 & 1 \end{pmatrix}$$

$$Y = \begin{pmatrix} \frac{\sqrt{2}}{2} & 0 & \frac{\sqrt{2}}{2} & 0 \\ 0 & 1 & 0 & 0 \\ -\frac{\sqrt{2}}{2} & 0 & \frac{\sqrt{2}}{2} & 0 \\ 0 & 0 & 0 & 1 \end{pmatrix}$$

$$Z = \begin{pmatrix} \cos\left(\arcsin\left(\sqrt{\frac{2}{3}}\right)\right) & -\sin\left(\arcsin\left(\sqrt{\frac{2}{3}}\right)\right) & 0 & 0 \\ \sin\left(\arcsin\left(\sqrt{\frac{2}{3}}\right)\right) & \cos\left(\arcsin\left(\sqrt{\frac{2}{3}}\right)\right) & 0 & 0 \\ 0 & 0 & 1 & 0 \\ 0 & 0 & 0 & 1 \end{pmatrix}$$

Multiplying the matrices from all these steps together, we have $K = MR_1T_1S$. For transforming from cell-type space $m^\mu$ to landscape space $\tilde{x}, \tilde{y}$, we define $H = K^{-1}$. $h_\mu^{x,y}$ denote the entries of $H$ such that $\tilde{x} = \sum_\mu h_\mu^x m^\mu$ and $\tilde{y} = \sum_\mu h_\mu^y m^\mu$.

#### *3. Correlated cell types and invariant transformations*

Cell types are often highly correlated, especially if they have common progenitors. Kanter and Sompolinsky [3] proposed an attractor network storage method which allowed for many highly-correlated attractors to coexist, which was further applied to the modern Hopfield network by Chaudhry et al. [4]. These generalized order parameters $m^\mu$ replace the usual magnetizations $m_\mu$. The new order parameters are defined as follows:

$$A_{\mu\nu} = \sum_i \xi_{\mu i}\xi_{\nu i}$$
$$m^\mu = \frac{1}{N}\sum_{\nu j}(A^{-1})^{\mu\nu}\xi_{\nu j}x_j = \sum_\nu (A^{-1})^{\mu\nu} m_\nu$$

These new order parameters are the result of a projection onto the non-orthogonal subspace of cell fates. It is a change of basis with $g_{\mu\nu} = A_{\mu\nu}$ acting as the metric tensor. We adopted Einstein notation to clarify when the metric is applied, with $g^{\mu\nu} = g^{-1}_{\mu\nu}$. The generalized order parameter is a contravariant vector defined with an upper index, $m^\mu = \sum_\mu g^{\mu\nu} m_\nu$, while the classic Hopfield order parameter is a covariant vector with a lower index $m_\mu$. The potential $V$ is a scalar, and any dot products used to calculate this scalar must take the metric tensor into account. For example, $V = \vec{m}\cdot\vec{m}$ would become $V = \sum_\mu m^\mu m_\mu = \sum_{\mu\nu} m_\mu g^{\mu\nu} m_\nu$.

The conversion from $(\tilde{x}, \tilde{y})$ coordinates to $(m^0, m^1, m^2)$ coordinates is defined in section A 2 as $\tilde{x} = \sum_\mu h^x_\mu m^\mu$ and $\tilde{y} = \sum_\mu h^y_\mu m^\mu$. Using the metric tensor, these are $\tilde{x} = \sum_{\mu\nu} h^x_\mu g^{\mu\nu} m_\nu$ and $\tilde{y} = \sum_{\mu\nu} h^y_\mu g^{\mu\nu} m_\nu$. Thus, the derivative $\frac{\partial V}{\partial m_\mu}$ involves the metric tensor as follows:

$$\frac{\partial V}{\partial m_\mu} = \frac{\partial V}{\partial \tilde{x}}\frac{\partial \tilde{x}}{\partial m_\mu} + \frac{\partial V}{\partial \tilde{y}}\frac{\partial \tilde{y}}{\partial m_\mu}$$
$$= \frac{\partial V}{\partial \tilde{x}}(\sum_\lambda h^x_\lambda g^{\lambda\mu}) + \frac{\partial V}{\partial \tilde{y}}(\sum_\lambda h^y_\lambda g^{\lambda\mu})$$

For clarity, since $h^{x,y}$ are actually 4-dimensional vectors, the notation used above corresponds to the following: in the products $\sum_{\mu\nu} h^{x,y}_\mu g^{\mu\nu} m_\mu$ and $\sum_\lambda h^{x,y}_\lambda g^{\lambda\mu}$, $g^{\mu\nu}$ and $m^\mu$ are moved to 4-dimensional space with the addition of a homogenous coordinate. These calculations are written out explicitly below.

$$\sum_{\mu\nu} h^{x,y}_\mu g^{\mu\nu} m_\nu \to h^{x,y}\cdot g\cdot m = \begin{pmatrix} h^{x,y}_0 \\ h^{x,y}_1 \\ h^{x,y}_2 \\ h^{x,y}_3 \end{pmatrix} \begin{pmatrix} g^{00} & g^{01} & g^{02} & 0 \\ g^{10} & g^{11} & g^{12} & 0 \\ g^{20} & g^{21} & g^{22} & 0 \\ 0 & 0 & 0 & 1 \end{pmatrix} \begin{pmatrix} m^0 \\ m^1 \\ m^2 \\ 1 \end{pmatrix}$$

The derivative $\frac{\partial V}{m_\mu}$ contains the terms $\sum_\lambda h^{x,y}_\lambda g^{\lambda\mu}$. In this case, $g^{\lambda\mu}$ is a 4-by-4 matrix as written above for the product, and after the product is calculated the fourth dimension is truncated so that the index $\mu$ refers once again to cell fates $0, 1, 2$. We can write this explicitly using $\alpha$ and $\beta$ to refer to the 4-dimensional space (three cell fate dimensions plus the extra dimension for transforming coordinates):

$$\sum_\lambda h^{x,y}_\lambda g^{\lambda\mu} \to \sum_\alpha h^{x,y}_\alpha g^{\alpha\beta} = (h^{x,y}\cdot g)_\beta$$

To bring this back to three coordinates for $\frac{\partial V}{\partial m_\mu}$, we simply take $\beta = 0, 1, 2$ and ignore $\beta = 3$.

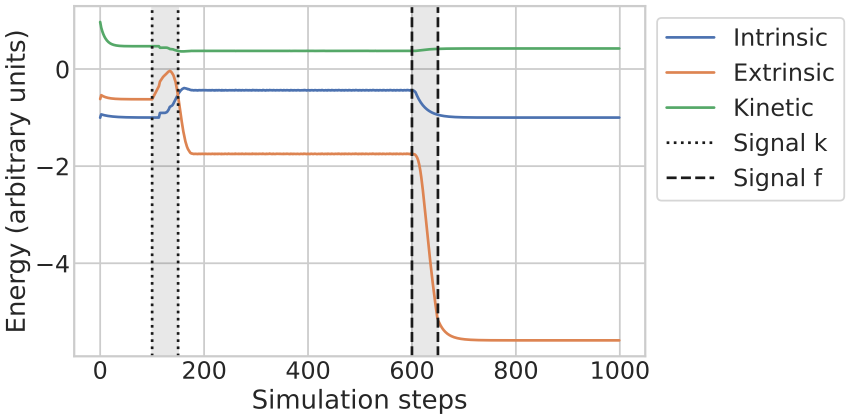


FIG. 5: The Hopfield energy and the imposed potential change as the simulation progresses. The imposed potential decreases while the Hopfield energy has a temporary increase. "Intrinsic" refers to the second term of $E$, the Hopfield energy. "Extrinsic" refers to the imposed potential $V$. "Kinetic" refers to the first term of the Hopfield energy $E$ . The units are arbitrary because, since the actual form of the Lyapunov function of the modified Hopfield model is not known, the scale has no meaning.

##### *4. Potentials over time*

It's well-known that the Hopfield model has an associated energy landscape which acts as the Lyapunov function of the system [5]. In other words, the update rule of the Hopfield model causes this energy function to decrease. For the modern Hopfield model, the energy takes the following form:

$$E = \frac{1}{2N}\sum_{i=1}^{N} x_i^2 - \log\left(\sum_{\mu}\exp(\beta m^{\mu})\right)$$

It's unclear what the corresponding Lyapunov function would be for the modified Hopfield model we introduce in this paper. However, we can compare how the intrinsic energy (defined above) and the extrinsic energy (the potential $V$ we insert into the system) change over time. We refer to the first term in the intrinsic energy as the "kinetic" term and the second term as the "intrinsic" term.

In Figure 5, we show how these different energies change over simulation time for the triple cusp landscape. This is the same simulation as shown at the top of Figure B 4: the first change in signalling (signal $f$) causes the cell to leave the first attractor basin, and the second change in signalling (signal $k$) causes the cell to leave the intermediate state and enter the final basin corresponding to a mature cell type. As shown in the figure, although the intrinsic energy increases temporarily when the cell enters a state not corresponding to one of the Hopfield stored patterns, the extrinsic energy decreases throughout the simulation.

#### B. Simulation details

The full details of the simulation can be found in the accompanying Python Jupyter notebooks. This section describes the structure of the code as well as some important aspects of the simulations performed to produce the figures in the main text.

Simulating a cell fate transition begins by instantiating an object of the `AttractorNetwork` class. The class simulates dynamics for the case of three primary attractor states. It works by storing the state of the network, $x_i$, and simulating the dynamics $\frac{dx_i}{dt}$ for specified landscape, signals, and attractor states. This class is structured to be modular, so it can take any suitable landscape as input. It requires the gradient of the landscape ($\frac{dV}{d\tilde{x}}, \frac{dV}{d\tilde{y}}$) and the attractor coordinates $\tilde{a}_0, \tilde{a}_1, \tilde{a}_2$ in $(\tilde{x}, \tilde{y})$ space. The gradient of the landscape is a function of the signaling parameters $(k, f)$, which control the bifurcations. The initial and final values of the signaling parameters $(k_{i,f}, f_{i,f})$ and the signal timing of the signals are specified at the start of the simulation. It uses this information, along with the attractor states $\xi_{\mu i}$, to calculate $h_\lambda^{x,y}$ and $g^{\lambda\mu}$, which are then used to calculate the update step:

$$x_i(t+1) = x_i(t) + \frac{dx_i}{dt}$$
$$\tau\frac{dx_i}{dt} = \sum_\mu \xi_{\mu i}\sigma^\mu(m^\mu(t) - \frac{\partial V}{\partial \tilde{x}}(\sum_\lambda h_\lambda^x g^{\lambda\mu}) - \frac{\partial V}{\partial \tilde{y}}(\sum_\lambda h_\lambda^y g^{\lambda\mu})) - x_i(t)$$

$\tau$ sets the speed of the dynamics. The result of varying $\tau$ or setting an asynchronous update schedule are discussed in Section B 4. The signal timing is discussed in Section B 1.

The simulations were run in gene expression space $(x_i)$ but the plots are shown are in cell fate space $(m_\mu)$. The cell fate coordinates were calculated using the attractor states $\xi_{\mu i}$. For figures 3, 2, the stored patterns used in the simulation were three scRNA-seq samples of alveolar type 1, alveolar type 2, and early epithelium cells. The former two are from Herriges et al. [6] while the latter is from Negretti et al. [7]. 17,970 genes were simulated.

#### *1. Signaling dynamics*

For Figure 2, the potentials were plotted in Mathematica using the equations shown. The values for the signaling parameters $(k, f)$ were varied as follows. For the landscape with a triple cusp, the system starts with $k = 0, f = 0$. The signaling parameter $k$ is increased from 0 to 0.3, destabilizing the attractor in which the system started. Then, $f$ is changed from 0 to 0.3, causing the intermediate attractor to disappear and tilting the landscape towards one of the final fates. In the landscape with a double cusp bifurcation, the parameters start at $k = 0.15, f = 0$. Parameter $k$ is kept at 0.15, and $f$ is changed from 0 to 0.1. This destabilizes the initial attractor and pushes the cell towards one of the final cell fates. The landscape with a heteroclinic flip bifurcation starts with $k = 0.5, f = 0$. Signal $k$ is shifted from 0.5 to 2 to destabilize the initial attractor, and before that bifurcation is fully complete, $f$ is changed from 0 to 0.5 to tilt the landscape towards the cell fate on the right-hand side.

In the simulated trajectories shown in Figures 3 and 2, the signaling parameters $f, k$ were varied manually in short time windows to reflect changes in the landscape caused by received signals. 10 trials were run for each landscape class. Not many trials were needed since the dynamics are so consistent (besides the added noise, see Section B 3) and there were enough time steps to produce a lot of data. Generally, the timing of signals was chosen to emphasize the features unique to each class. If the signals are changed very quickly, the system quickly converges to its final point. If the signals are changed slowly enough, one can find cells along the entire trajectory.

Start and end times, as well as start and end values, are specified for each parameter when the `AttractorNetwork` class is instantiated. $t_s^f, t_e^f$ are the start and end times for parameter $f$ while $t_s^k, t_e^k$ are the start and end times for parameter $k$. $(f_s, f_e), (k_s, k_e)$ indicate the start and end values for each of the parameters. Within the time window specified by the start and end times for each signal, the parameter was changed as follows:

$$f(t+1) = f(t) + \frac{f_e - f_s}{t_e^f - t_s^f}$$
$$k(t+1) = k(t) + \frac{k_e - k_s}{t_e^k - t_s^k}$$

For the triple cusp, the two signals represent two stages of differentiation. The first stage is the transition from the initial state to the intermediate state, and the second stage is the specification into one of the final two fates. Each simulation was run for 1000 time steps. The signaling schedule was chosen so that the cell transitioned to the intermediate attractor first, before receiving signals that push it to its final fate. The initial signaling parameters were $k_s = 0, f_s = 0$. From time steps 0 to 400, $k$ was gradually changed from $k_s = 0$ to $k_e = 0.3$. This change represents an initial signal that pushes the cell out of its initial attractor state. Then, from steps 600 to 700, $f$ was changed from $f_s = 0$ to $f_e = \pm 0.3$. This represents a signal that pushes the cell to one of the final two states. The sign of the signal is chosen randomly for each trial in order to show routes to both final fates. The first signal was varied over a longer stretch of time than the second to show cells along the first leg of the path. Also, there is a time gap between the signals of 200 time steps. The length of this gap determines how long cells stay in the intermediate attractor and therefore how prominent that cluster is in the plots. If the time between signals is too short, the intermediate attractor is difficult to detect.

In the double cusp, changing either signal can destabilize the attractor. $f$ represents the bias towards one of the two final fates. If $k$ is changed while $f$ remains zero, the initial attractor basin becomes a saddle point, and the cell has equal probability to end up in either final attractor if it is perfectly positioned on the saddle point. The bifurcation from changing $f$ is more interesting than the one from changing $k$ because the latter only involves a random choice rather than a decision, so we only varied $f$ for this bifurcation. The simulation ran for 750 time steps per trial. At the beginning of the simulation, $f = 0$ and $k = 0.15$. For the duration of the entire simulation, $f$ is gradually changed to $\pm 0.1$. As for the triple cusp, the sign of this signaling parameter was randomly chosen for each trial in order to cover both possible paths in the transition.

In the heteroclinic flip landscape, the first signal pushes the cell out of the initial attractor towards a saddle point and the second signal biases the cell towards one of the final two fates. In the simulations, this system starts with $f_s = 0, k_s = 0.5$. Changing $k$ destabilizes the initial attractor. Once the initial attractor is destabilized, the system is very sensitive to changes in $f$ because it is poised at a saddle point between the final two fates. For this class, we chose to have an overlap in the time windows of signals. This is to avoid a situation where the cell moves out of the initial attractor towards the saddle point, starts moving towards one of the final fates and then switches directions due to a change in $f$. Each simulation was run for 750 steps. From steps 100 to 500, $k$ was changed from $k_s = 0.5$ to $k_e = 2$. From steps 450 to 600, $f$ was changed from $f_s = 0$ to $f_e = \pm 0.5$. Similarly to the other two landscapes, the sign of $f_e$ was chosen randomly for each trial in order to show trajectories to both final fates.

#### *2. Inverse temperature*

For all simulations except when explicitly stated otherwise, the inverse temperature $\beta$ in the softmax of the update rule was set to $\beta = 2N$, where $N$ is the number of genes simulated. For the modern Hopfield network to converge to the correct pattern, it is necessary that the temperature is low enough such that $e^{\beta} \gg P$, where $P$ is the number of stored patterns. This can be seen by considering the softmax in the update rule when $m^{\mu} = 1, m^{\nu \neq \mu} = 0$:

$$\begin{aligned}
\sigma(\beta m_{\mu}) &= \frac{e^{\beta}}{e^{\beta} + e^{0} + e^{0} + ...} \\
&= \frac{1}{1 + Pe^{-\beta}} \\
\sigma(\beta m_{\nu \neq \mu}) &= \frac{e^{0}}{e^{\beta} + e^{0} + e^{0} + ...} \\
&= \frac{1}{e^{\beta} + P}
\end{aligned}$$

In this case, the update rule should keep the state static, so $\sigma(\beta m^{\mu}) = 1$ and $\sigma(\beta m^{\nu \neq \mu}) = 0$. This condition is met for $e^{\beta} \gg P$.

#### *3. Noise*

Noise was added to the simulation in Figures 2 and 3 to imitate the noise in scRNA-seq and facilitate comparison between simulated trajectories and data. The simulations were initiated at the initial attractor state with noise added to the gene expression: $x_i(t = 0) = \xi_{0i} + z_i$. The noise $z_i$ for each gene $i$ is drawn from identical, independent normal distributions. Modern Hopfield networks are excellent at converging and de-noising, so after one step the simulation converges to the noise-less trajectory. To maintain noise along the path, we added noise to the recorded variables. Specifically, normally-distributed noise was added to the observed $x_i$ at each step, and a fraction of counts were randomly set to zero to imitate the high levels of dropout in scRNA-seq [8]. Additionally, a large percent of the simulated cells have not been included in the plot, to reflect the fact that scRNA-seq experiments can't capture every cell at every step of the process. The addition of noise makes it harder to distinguish between classes, but contour plots (as in Figure 3) help highlight important features.

#### *4. Robustness to update schedule*

In our model, all genes are updated simultaneously. However, this does not reflect biological reality. To reflect the asynchronous nature of gene regulation, we can randomly update some genes at each simulation step and not others. The model is robust to such changes, and still converges to the expected basins. This is illustrated in Figure B 4, where a different percentage of genes are randomly chosen to be updated in the simulation of a triple cusp bifurcation. The only difference caused by this change in update schedule is a longer time to converge to the attractor basin. This timing only matters relative to the timing of signaling; as long as it occurs on faster timescales than signaling, the basic behavior of the system is not affected.

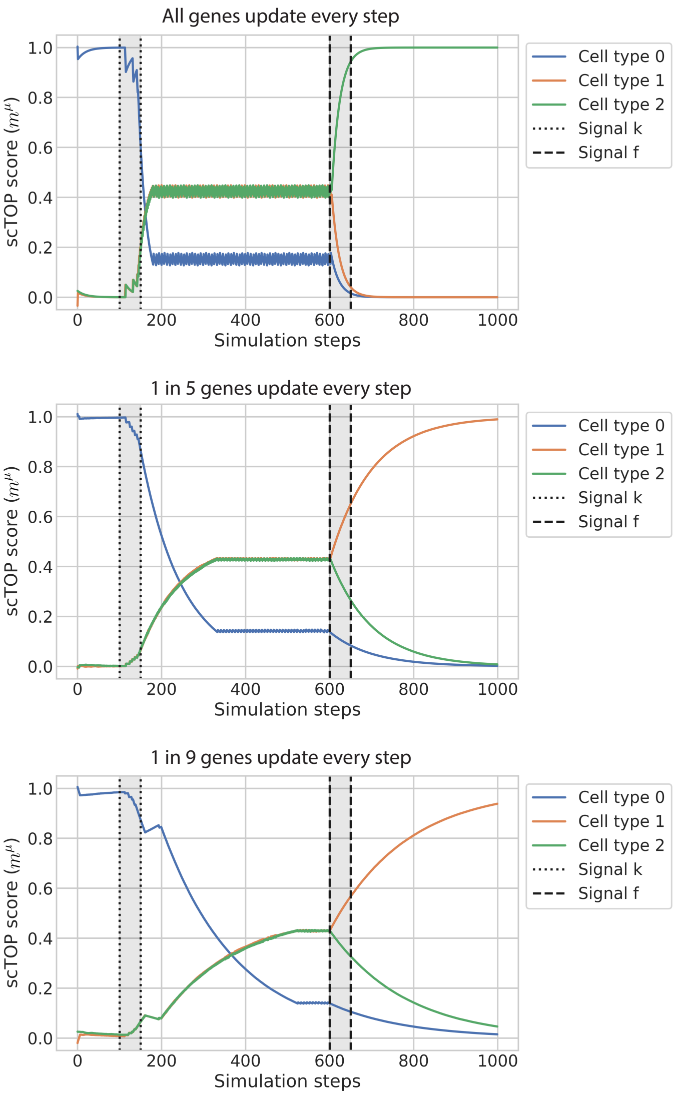


FIG. 6: The dynamical system is robust to changes in update schedule. All three plots show the change in cell type alignment in a triple cusp bifurcation over the course of simulation time, where the changes in signal $\Delta f$ and $\Delta k$ occur in the indicated intervals. In each plots, the fraction of genes randomly chosen to be updated at each simulation step is varied, ranging from all genes (top) to 1/5 genes (middle) to 1/9 genes (bottom). The basic behavior remains the same regardless of how many genes are updated, although the time scales vary.

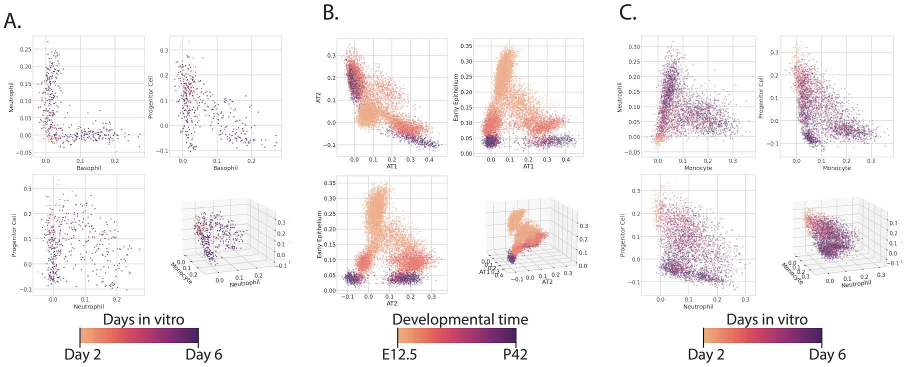


FIG. 7: The same data as in Figure 4, shown as a scatter plot colored by time point. The axes correspond to the two mature fates relevant to each experiment as well as a progenitor cell type, to show the maturation from an early state to a later one.

#### C. Limitations of the model

Because decision-making classes are describe general characteristics, there are many facets of our simulations that were not constrained by choice of bifurcation. In the future, with more analysis using scTOP scores to plot data on the order parameter axes, it may be possible to place more constraints on the simulations. For this paper, we tried to make choices that minimized additional assumptions. For example, in transforming between coordinates, we chose to align the attractor coordinates in $x - y$ space with the simplex in $\vec{m}$ space.

The paths of transition predicted by our model are highly dependent on the choice of potential, control parameters, and other mathematical details. Although the decision-making classes are generic, the choices we made in choosing the forms of the potentials and in transforming between the 2-dimensional $V(x, y)$ and the three-dimensional $V(m^0, m^1, m^2)$ are not generic. With the changes of control parameters, the attractor basins shift slightly in position, which is not necessarily reflective of actual observations. With more scTOP analyses, it will be more and more possible to fit these various parameters and make more informed decisions.

Another limitation is the fact that it is not possible to fully determine the landscape. This is an inherent aspect of coarse-grained models. Not only are there infinite possibilities for connecting decision-making classes, but different landscapes can produce the exact same trajectories in certain regimes (see section A 1). It is easier for the model to eliminate particular classes of bifurcations than definitively identify the exact class.

---
